# Enhanced activity of glycolytic enzymes in *Drosophila* and human cell models of Parkinson’s disease based on *DJ-1* deficiency

**DOI:** 10.1101/2020.03.10.985135

**Authors:** Cristina Solana-Manrique, Francisco José Sanz, Edna Ripollés, M. Carmen Bañó, Josema Torres, Verónica Muñoz-Soriano, Nuria Paricio

## Abstract

Parkinson’s disease (PD) is a neurodenerative debilitating disorder characterized by progressive disturbances in motor, autonomic and psychiatric functions. The pathological hallmark of PD is the loss of dopaminergic neurons in the substantia nigra pars compacta, which causes striatal dopamine deficiency. Although most PD cases are sporadic (iPD), approximately 5-10% of all patients suffer from monogenic PD forms caused by highly penetrant rare mutations segregating with the disease in families (fPD). One of the genes linked to monogenic PD is *DJ-1*. Mutations in *DJ-1* cause autosomal recessive early-onset forms of fPD; however, it has been shown that an over-oxidized and inactive form of the DJ-1 protein is found in the brains of iPD individuals. Valuable insights into potential PD pathogenic mechanisms involving DJ-1 have been obtained from studies in cell and animal PD models based on *DJ-1* deficiency such as *Drosophila*. Flies mutant for the *DJ-1β* gene, the *Drosophila* ortholog of human *DJ-1*, exhibited disease-related phenotypes such as motor defects, increased reactive oxygen species production and high levels of protein carbonylation. In the present study, we show that loss of *DJ-1β* function significantly increased the activities of several regulatory glycolytic enzymes. Similar results were obtained in *DJ-1*-deficient SH-SY5Y neuroblastoma cells, thus suggesting that loss of *DJ-1* function in both PD models produces an enhancement of glycolysis. Our results also show that FDA-approved compounds such as meclizine and dimethyl fumarate, which have different clinical applications, are able to attenuate PD-related phenotypes in both models. Moreover, we found that they could exert their beneficial effect by increasing glycolysis through the activation of key glycolytic enzymes. Taken together, these results are consistent with the idea that increasing glycolysis could be a potential disease-modifying strategy for PD, as recently suggested. Besides, they also support further evaluation and potential repurposing of meclizine and dimethyl fumarate as modulators of energy metabolism for neuroprotection in PD.

## INTRODUCTION

Parkinson’s disease (PD) is a neurodenerative debilitating disorder characterized by progressive disturbances in motor, autonomic and psychiatric functions. The pathological hallmark of PD is the loss of dopaminergic (DA) neurons in the substantia nigra pars compacta, which causes striatal dopamine deficiency. In some cases, nigral neuronal loss is accompanied by the presence of proteinaceous aggregates in the remaining neurons, the so-called Lewy bodies [1,2]. The exact disease mechanisms underlying DA neuron loss, which eventually lead to clinical PD, are still unclear and probably involve a combination of genetic predispositions and environmental factors as well as lifestyle exposures. Indeed, multiple pathways and mechanisms appear to be implicated in PD pathogenesis. Thus, accumulation of misfolded proteins aggregates, mitochondrial dysfunction, oxidative stress (OS), energy failure, neuroinflammation or genetic mutations have been proposed to contribute to the onset and progression of PD [1,3,4]. Although highly effective therapies are available to relieve PD symptoms, mainly focused on reconstituting DA signaling in the surviving neurons, none of these treatments is curative since they are unable to restore the lost/degenerated DA neurons or to prevent or delay the disease progression [2,3].

Most PD cases are idiopathic; however, approximately 5-10% of all patients suffer from monogenic PD forms caused by highly penetrant rare mutations segregating with the disease in families. Those familial PD (fPD) cases are, very often, clinically indistinguishable from idiopathic PD (iPD) except for age at onset [5]. Several genes whose mutations cause PD have been identified in the last two decades. Studying those genes, their function as well as the pathways they are involved in has been crucial to discover most of the currently known mechanisms underlying the neuropathology of the disease [1,2,6]. One of the genes unequivocally linked to monogenic PD is *DJ-1*. Mutations in *DJ-1* cause autosomal recessive early-onset forms of fPD [7]; interestingly, it has been shown that an over-oxidized and inactive form of the DJ-1 protein is found in the brains of iPD individuals [8]. Although the *DJ-1* gene was initially identified as an oncogene, a number of experimental results suggested that it is also related to onset or physiology of several OS-induced diseases, including stroke, type 2 diabetes mellitus, male infertility, and neurodegenerative diseases other than PD (see [9]). Consistently, DJ-1 appears to be a multitask protein shown to perform a variety of functions including transcriptional regulation of antioxidant genes, quenching reactive oxygen species (ROS), regulation of signal transduction pathways, and roles as chaperone and protease [9]. Valuable insights into potential PD pathogenic mechanisms involving DJ-1 have been obtained from studies in cell and animal PD models based on *DJ-1* deficiency such as those developed in *Drosophila* (reviewed in [10]). Indeed, flies mutant for the *DJ-1β* gene, the *Drosophila* ortholog of human *DJ-1*, exhibit disease-related phenotypes such as motor defects and increased reactive oxygen species (ROS) levels [11–13]. Moreover, high levels of protein carbonylation, a post-translational oxidative modification that proteins suffer in high OS conditions were observed in PD model flies [13]. Oxidative damage to proteins may alter expression levels and/or confer a toxic loss or gain of function [14]. Thus, identifying proteins specifically carbonylated in cell/animal PD models may lead to discover pathogenic mechanisms and to identify new targets to alter the course of the disease [15].

Several studies have revealed links between glucose metabolism and alterations in cellular bioenergetics, redox homeostasis and cell death induced by PD-related risk factors. While a decrease in glucose metabolism and abnormally levels of lactate/pyruvate have been observed in PD patients, other studies in different PD models have demonstrated that glycolysis is upregulated in response to mitochondrial dysfunction (reviewed in [16]). This upregulation may counteract energy failure and DA cell death occurring when mitochondrial function is impaired [17]. Accordingly, it was reported that mouse *DJ-1*-null embryonic fibroblasts (MEFs) showed a higher glycolytic flux than wild-type MEFs [18]. The authors proposed that DJ-1 represses glycolysis and cell proliferation by transcriptionally activating *Pink1*, another PD-related gene whose loss was also reported to up-regulate glucose metabolism in order to sustain cell proliferation [19].

In such a scenario, we hypothesized that a similar enhancement in glycolysis could be occurring in fly and human cell PD models based on *DJ-1* inactivation. Preliminary results obtained from a redox proteomics assay performed in *DJ-1β* mutant flies led us to identify two glycolytic enzymes, enolase and phosphofructokinase, whose carbonylation levels were increased in PD model flies compared to controls (C.S.-M., F.J.S., V.M.-S. and N.P., in preparation), thus suggesting that the glycolytic pathway might be altered in such flies. In the present study, we show that loss of *DJ-1* function significantly increased the activities of these and other key regulatory glycolytic enzymes like hexokinase and pyruvate kinase in flies and human neuroblastoma cells. These results supported the hypothesis that glycolysis may be enhanced in both PD models. Moreover, our results also revealed that a further increase of the glycolytic pathway by administration of compounds such as meclizine and dimethyl fumarate had potential disease-modifying effects in fly and cell PD models. These results are consistent with recent findings in which increasing glycolysis with compounds like terazosin, the tetramethylpyrazine analogue T-006 or hydrogen sulfide was shown to attenuate disease-related phenotypes in different experimental PD models [20–22], thus pointing to the glycolytic pathway as a potential therapeutic target for PD.

## MATERIALS AND METHODS

### *Drosophila* strains

All stocks and crosses were cultured on standard *Drosophila* food at 25 °C, unless otherwise indicated. The stocks used in this work were: *y,w* (<u>B</u>loomington <u>D</u>rosophila <u>S</u>tock <u>C</u>enter #6598: *y*^*1*^,*w*^*1118*^), *w; DJ-1β*^*ex54*^ (hereafter called *DJ-1β;* [23]), *arm-GAL4* (BDSC #1560: *P{GAL4-arm.S}11*), *UAS-iRPfk* (Vienna Drosophila Research Center #3017: *w*^*1118*^; *P{GD1508}v3017*), and *UAS-iREno* (BDSC #26300: *y*^*1*^ *v*^*1*^; *P{TRiP.JF02070}attP2*). The *y,w* line was used as wild-type control in experiments with the *DJ-1β* strain. For RNAi experiments, the *arm-GAL4* driver line was crossed with UAS-*iRPfk* and UAS*-iREno* lines to ubiquitously downregulate *Pfk* and *Eno* expression, respectively. Progeny of these crosses is referred to as *iRPfk* and *iREno*, respectively. In these experiments, progeny of crosses between *y,w* and *arm-GAL4* flies was used as control.

### Cell culture and drug treatment

SH-SY5Y neuroblastoma control cells (*pLKO.1*) and *DJ-1-*deficient cells generated in our laboratory (Sanz et al. 2017) were maintained in a selective growth medium consisting of Dulbecco’s Modified Eagle Medium/Nutrient Mixture F-12 (DMEM/F-12) (Labclinics) supplemented with 2 μg/ml puromycin (Labclinics), 10% (v/v) fetal bovine serum (FBS) (Labclinics) and 100 mg/ml penicillin/streptomycin (Labclinics) at 37 °C and 5% CO_2_. Cell viability assays with different concentrations of dimethyl fumarate (DMF) and meclizine (MEC) were performed in *DJ-1*-deficient cells with MTT (3-(4, 5-dimethylthiazol-2-yl)−2–5-diphenyltetrazolium bromide) (Sigma-Aldrich) as described in [24], with slight modifications. Briefly, *DJ-1-*deficient cells were seeded in a 96-well plate at a density of 1.8×10^4^ cells/well and incubated for 24 h with different concentrations of DMF and MEC in the range 1-50 µM, or with 0.1% dimethyl sulfoxide (DMSO) as vehicle. Subsequently they were incubated with 100 μM H_2_O_2_ for 3 h. As a result of these experiments, the most effective concentration for each compound in suppressing OS-induced cell death was selected for further analysis.

### Protein extracts

Enzymatic activities were measured in protein extracts from 15-day-old *DJ-1β* mutants or control flies either untreated or treated with DMF or MEC. In all cases, twenty female flies were homogenized in 200 μl of 50 mM Tris-HCl buffer, pH 7.4 with a steel bead in a TissueLyser LT (Qiagen) for 5 min at 50 Hz. Fly extracts were centrifuged at 9000*g* for 8 min at 4°C. After centrifugation supernatant was collected and Halt Protease Inhibitor Cocktail (ThermoScientific) added.

Enzymatic activities were also determined in protein extracts from *DJ-1-*deficient and control (*pLKO.1*) SH-SY5Y cells in which OS was induced by incubation with 50 μM H_2_O_2_. Extracts were obtained from aliquots of 3×10^6^ cells of mutant and control cells either untreated or pretreated for 24 h with DMF or MEC. To do this, cells were lifted with trypsin and harvested in complete DMEM-F12 medium, centrifuged at 300g for 5 min, washed with 1% PBS buffer at 4 °C and resuspended in FT lysis buffer [200 mM Tris-HCl pH 7.8, 600 mM KCl, 20%glycerol (v/v)]. Cell suspensions were subjected to three successive cycles of freezing in liquid N_2_, thawing on ice and vortexing. Lysates were centrifuged at 15.000g for 10 min at 4 °C and Halt Protease Inhibitor Cocktail (ThermoScientific) was added. Supernatants containing fly and cell lysates were quantified with BCA Protein Assay Kit (Pierce) following manufacturer’s instructions.

### Enolase and phosphofructokinase activity assays

25 μg of protein extracts for enolase (Eno; EC 4.2.1.11) and 40 μg of protein extracts for phosphofructokinase (Pfk; EC 2.7.1.11) were added to 100 µl of the assay buffers in a 96-well plate. Compositions of buffers depending on the enzymatic activity to be assayed were as follows: <u>Eno</u>: 100 mM KCl, 10 mM MgCl_2_, 20 mM HEPES pH 7, 1.75 mM ADP, 0.5 mM NADH, 15 U/ml pyruvate kinase, 15 U/ml lactate dehydrogenase, pH 7.4; <u>Pfk</u>: 91.1 mM Tris-HCl pH 9, 1 mM ATP, 5 mM KCl, 2 mM MgSO_4_, 0.728 mM phosphoenolpyruvate, 0.5 mM NADH, 5 U/ml pyruvate kinase, 10 U/ml lactate dehydrogenase, pH 8. Proteins were incubated with assay buffer for 5 min to test for possible absorbance variations and, after this time, reaction was started by adding 2 mM phosphoglycerate for Eno and 10 mM fructose 6-phosphate for Pfk. Absorbance was measured at 340 nm using an Infinite 200 PRO reader (Tecan) every 30 s for 20 min at 25°C for fly extracts and 37°C for cell extracts. In all cases sample absorbance levels were measured, subtracting their corresponding blanks. Enzymatic activity was calculated from the decrease in absorbance at 340 nm due to NADH to NAD^+^ oxidation. The experiments were validated in *iReno* and *iRpfk* mutant flies, in which the expression of the genes encoding the corresponding enzymes is reduced (Fig. S1B). All the assays were performed in triplicate.

### Hexokinase and pyruvate kinase activity assays

To measure these enzymatic activities we used assays based on previously published methods performed in *Drosophila* [25]. In these assays, 15 μg of protein extracts were added to 100 µl of the assay buffers for pyruvate kinase (PyK in *Drosophila* or Pk in human cells; EC 2.7.1.40) and hexokinase (Hex-A in *Drosophila* or Hk in human cells; EC 2.7.1.1) in a 96-well plate. Compositions of buffers depending on the enzymatic activity to be assayed were as follows: <u>Hex-A or Hk</u>: 20 mM Tris-HCl pH 9, 2 mM MgCl_2_, 8 mM ATP, 100 mM glucose, 1 mM NADP^+^, 5 U/ml glucose 6-phosphate dehydrogenase, 10 U/ml lactate dehydrogenase, pH 7.3; <u>PyK or Pk</u>: 50 mM HEPES pH 7.4, 60 mM KCl, 8 mM MgSO_4_, 2 mM ADP, 2 mM phosphoenolpyruvate, 0.5 mM NADH, 10 U/ml lactate dehydrogenase, pH 7.3. The absorbance was measured at 340 nm using an Infinite 200 PRO reader (Tecan) for 10 min at 25°C for fly extracts and 37°C for cell extracts. In all cases sample absorbance levels were measured, subtracting their corresponding blanks. PyK or Pk activity was calculated from the decrease in absorbance at 340 nm due to NADH to NAD^+^ oxidation. Hex-A or Hk activity was calculated from the increase in absorbance at 340 nm due to NADP^+^ to NADPH reduction. All the assays were performed in triplicate.

### RT-qPCR analyses

Total RNA from ten 15-day-old *DJ-1β, iRPfk, iREno* or control flies (*y,w* or *arm*-GAL4/+, depending on the experiment) and from aliquots of 3×10^6^ *DJ-1* mutant and control cells was extracted with TRItidy G™ (Panreac, AppliChem) following manufacturer’s instructions. RNA was reversed transcribed using *FIREScript RT cDNA Synthesis MIX* (Solis BioDyne). Quantitative real-time RT-PCR (RT-qPCR) was performed using EvaGreen (Solis BioDyne) on an ABI StepOnePlus™ Real-Time PCR System (Applied Biosystems). *tubulin* levels were measured and used as an internal control for RNA amount in each sample. Data analysis was performed in quadruplicate experiments with StepOnePlus™ software v2.3. Results are expressed as relative expression of the genes in mutant flies or cells compared to controls.

For the RT-qPCR reactions, we used the following pairs of primers: *Pfk* direct primer (5’-CGACCTCATTGCAGAGACGA-3’); *Pfk* reverse primer (5’-ACCACTGCTTCTTCGGGATG-3’); *Eno* direct primer (5’-ACCAGCCCCATCGAATGAGA-3’); *Eno* reverse primer (5’-TCCAGCTCCCTCTCCAAGAT-3’); *Hex-A* direct primer (5’-CAAAATCAGCGACAGCGACC-3’); *Hex-A* reverse primer (5’-GCGGGCTGTGAGTTGTAAGA-3’); *PyK* direct primer (5’-TCTTGGTGACTGGCTGAAGG-3’); *PyK* reverse primer (5’-GCCGTTCTTCTTTCCGACCT-3’); *PFKM* direct primer (5’-CTGCCCCTCATGGAATGTGT-3’); *PFKM* reverse primer (5’-CCTCTCAGCTTCAGGGCTTC-3’); *PFKP* direct primer (5’-ATCATCGGTGGATTCGAGGC-3’); *PFKP* reverse primer (5’-TTGGACACAGTAGCGGGAAC-3’); *ENO1* direct primer (5’-ACCCAAAGAGGATCGCCAAG-3’); *ENO1* reverse primer (5’-AACCAGGTCAGCGATGAAGG-3’); *ENO2* direct primer (5’-TGCACAGGCCAGATCAAGAC-3’); *ENO2* reverse primer (5’-ACAGCACACTGGGATTACGG-3’); *HK1* direct primer (5’-CATGCTGCTGGAGGTGAAGA-3’); *HK1* reverse primer (5’-CAGGGCCAAGAAGTCACCAT-3’); *PKM* direct primer (5’-GGTCCTGGGAGAGAAGGGAA-3’); *PKM* reverse primer (5’-TAGATCACCACGAGCCACC-3’); *Tubulin Drosophila* direct primer (5’-GTATCTCTATCCATGTTGGTCAGG-3’); *Tubulin Drosophila* reverse primer (5’-AGACGGCATCTG GCCATCG-3’); *Tubulin Homo sapiens* direct primer (5’-GCCGAGATCACCAATGCCT-3’); *Tubulin Homo sapiens* reverse primer (5’-TCACACTTGACCATTTGATTGGC-3’).

### Drug feeding in *Drosophila*

Flies were cultured on standard *Drosophila* food containing 0.1% DMSO for untreated control experiments, or in the same medium supplemented with a final concentration of 50 µM MEC or 7 µM DMF for treatment experiments. For MEC, we used the highest concentration possible in which the final DMSO concentration was 0.1% (as in controls). Regarding DMF, 7 µM was shown to be the most effective concentration in *Drosophila* climbing assays (data not shown). Ten males and 20 females of *y,w* or *DJ-1β* flies were crossed in each vial with or without compound and maintained at 25 °C. For climbing assays, freshly eclosed flies were transferred to new vials with the compounds and the climbing assay was carried out 5 days after. For enzymatic assays, freshly eclosed flies were transferred to new vials with or without the compounds. Vials were replaced every two days for 15 days. After this time, 15-day-old flies were frozen in liquid N_2_ and stored at -80 °C.

### Climbing assay

Locomotor ability of flies treated with compounds or DMSO as vehicle medium was analyzed with a climbing assay as previously described in [24]. The experiments were carried out three times and the climbing ability of control and mutant flies was determined as the average of the height reached by each fly after 10 s.

### Quantification of H_2_O_2_ levels

H_2_O_2_ levels were measured using the *Amplex H*_*2*_*O*_*2*_ *Red Kit* (Invitrogen) as described in [24]. Briefly, ten 5-day-old females were homogenized in 200 μl of 50 mM Tris–HCl, pH 7.4, with a steel bead in a TissueLysser LT (Qiagen) as described above. Fly extracts were transferred to a 96-well fluorescence plate (Greiner 96 well black plate, polypropylene). Fluorescence was monitored, according to manufacturer’s protocol, using an Infinite 200 PRO reader (Tecan). H_2_O_2_ levels are expressed as fluorescence intensity in arbitrary units per mg of protein. Data were relativized to values obtained in *DJ-1β* flies cultured in DMSO. All experiments were carried out using three biological replicates and three technical replicates for each sample.

### Determination of protein carbonyl group formation

Protein carbonyl groups were measured in 5-day-old fly extracts using 2,4-dinitrophenylhydrazine (DNPH) derivatization in 96-well plates (Greiner 96 well plate, polypropylene) as described in [26]. Briefly, 50 µl of 10 mM DNPH was added to 50 µl of fly extracts for 10 min. Next, 25 µl of 6 M NaOH was added. After 10 min of incubation, absorbance was read at 450 nm using an Infinite 200 PRO reader (Tecan). All experiments were carried out using three biological replicates and three technical replicates for each sample.

### Statistical analysis

Data are expressed as means ± standard deviation (s.d.). The significance of differences between means was assessed using t-test. Differences were considered significant when \**P* < 0.05.

## RESULTS

### Activity of glycolytic enzymes is increased in *DJ-1β* mutant flies

It is commonly known that OS plays an important role in the pathogenesis of PD [17,27]. One consequence of elevated ROS production is protein carbonylation, a posttranslational modification that may interfere in cell signaling and other cellular processes because it leads to protein degradation, inactivation or activation [14,28]. We previously demonstrated that the global protein carbonyl content was significantly higher in 15-day-old *DJ-1β* mutants than in control flies, consistent with increased ROS levels in those flies as a consequence of *DJ-1β* function loss [13]. Those results supported the hypothesis that protein oxidative damage due to high OS levels may represent an important pathogenic event in PD [24]. In order to investigate the molecular pathways that could be affected by *DJ-1β* deficiency as well as to identify putative PD biomarkers, we carried out a redox proteomic study in 15-day-old PD model and control flies. A total of 53 proteins showing differential carbonyl levels in *DJ-1β* mutant flies compared to *y,w* controls, but that did not show significant differences in expression levels, were detected (C.S.-M., F.J.S., V.M.-S. and N.P., in preparation). Among the proteins that exhibited increased carbonylation levels in mutant flies, we identified Pfk and Eno, both involved in glucose metabolism, thus suggesting that glycolysis may be altered in PD model flies. To confirm this assumption, we measured the activities of Pfk and Eno enzymes in *DJ-1β* mutant flies by designing specific assays that were first tested in protein extracts from *iRPfk* and *iREno* flies, in which the expression of the corresponding genes was ubiquitously downregulated with the *arm*-GAL4 driver. As expected, Pfk and Eno activities were significantly reduced in *iRPfk* and *iREno* flies compared to controls (Fig. S1A). qRT-PCR analyses in *iRPfk* and *iREno* mutants confirmed that *Pfk* and *Eno* gene expression, respectively, was decreased in these flies (Fig. S1B). Subsequently, we analyzed Pfk and Eno activities in 15-day-old *DJ-1β* mutants and *y,w* control flies using the same assays. Our results showed that both activities were increased in PD model flies compared to controls, especially Pfk (Fig. 1A). Therefore, our results suggested that glycolysis could be enhanced in *DJ-1β* mutants. At this point we decided to further confirm this finding by analyzing the activity of other glycolytic enzymes essential for the process. Glycolysis is a highly regulated pathway in which Pfk is the main regulatory enzyme; however, other enzymes like hexokinase (Hex-A) and pyruvate kinase (PyK) are also involved in important regulatory steps of this pathway [29]. Considering this, we analyzed the activity of Hex-A and PyK in protein extracts of 15-day-old *DJ-1β* mutants and control flies. To this end, we used published protocols that were already shown to efficiently measure the activity of these enzymes in *Drosophila* [25]. Our results showed that while PyK activity was highly increased in *DJ-1β* mutant flies compared to controls, no significant differences in Hex-A activity were observed (Fig. 1B). RT-qPCR analyses in PD model and control flies indicated that changes in the activities of the glycolytic enzymes in *DJ-1β* mutants were not reflecting changes in gene expression levels (Fig. S2). Hence, the fact that Pfk, Eno and PyK activities are increased in *DJ-1β* mutants strongly suggests that the glycolytic pathway is enhanced in this *Drosophila* PD model. Our results are consistent with those reported in a previous study performed in MEFs obtained from *DJ-1* mutant mice, in which an enhancement of the glycolytic rate was also observed [18].

**Figure 1.**
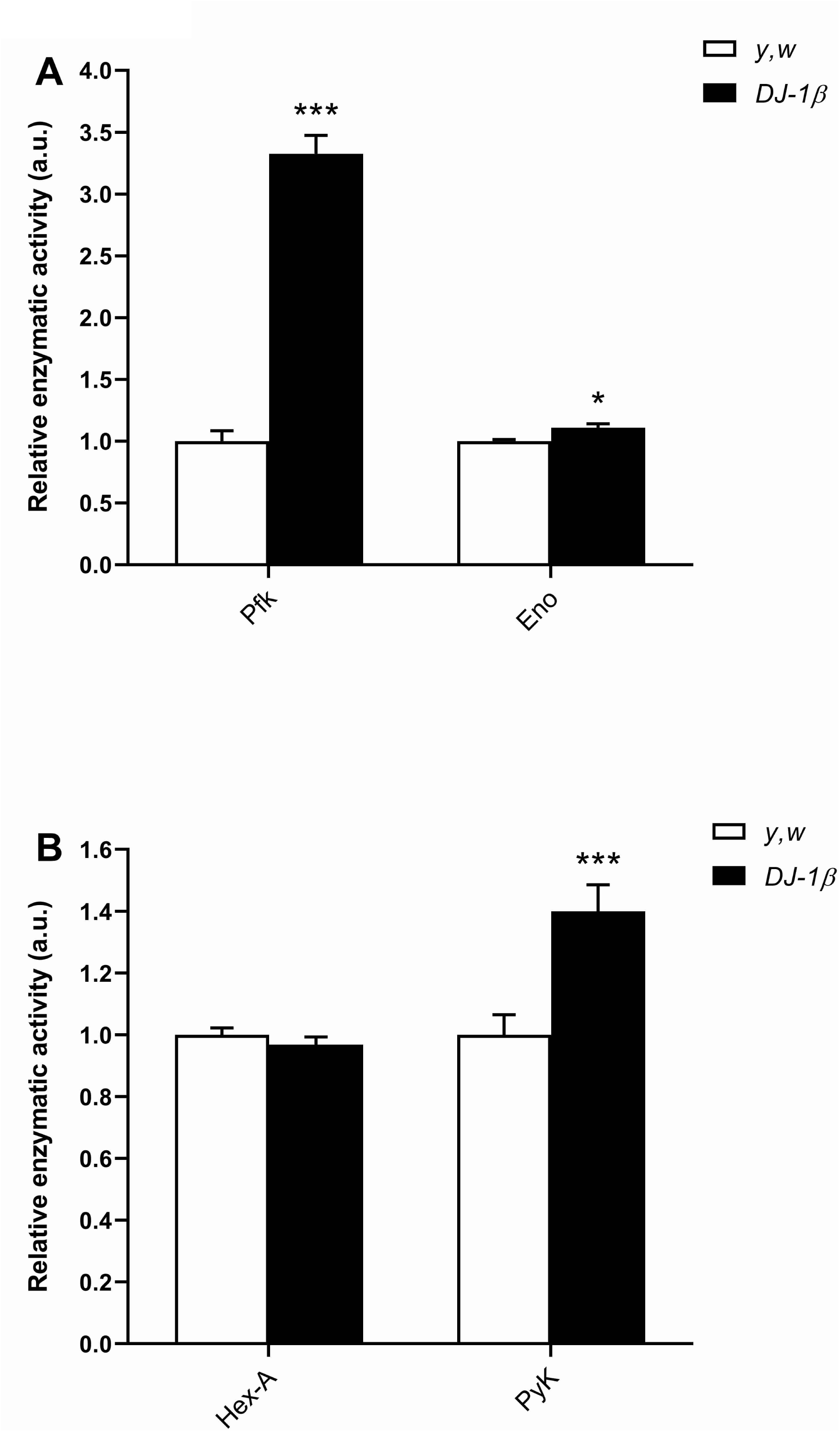
Activity of glycolytic enzymes in *DJ-1β* mutant flies. (A) Phosphofructokinase (Pfk) and enolase (Eno) enzymatic activities in 15-day-old *DJ-1β* mutant flies. (B) Hexokinase A (Hex-A) and pyruvate kinase (PyK) enzymatic activities in 15-day-old *DJ-1β* mutant flies. In all cases results were relativized to data obtained from 15-day-old *y,w* control flies and are expressed as arbitrary units (a.u.). Error bars show s.d. from three independent experiments in which three biological replicates were used (*, *P* < 0.05; ***, *P* < 0.001).

### Activity of glycolytic enzymes is increased in *DJ-1-*deficient human cells

To verify the relevance of our findings in *Drosophila*, in which enzymatic activities were analyzed in extracts from whole animals, we performed the same assays in an *in vitro* human cell PD model generated in neuron-like cells. To this end, the activities of the key regulatory glycolytic enzymes Pfk, Hk (hexokinase), Pk (pyruvate kinase) and of Eno were determined in human neuroblastoma SH-SY5Y cells mutant for the *DJ-1* gene [24]. We and others already showed that *DJ-1-*deficient SH-SY5Y neuroblastoma cells were more susceptible than control cells to OS-induced cell death [24,30,31]. Accordingly, dose-dependent citotoxicity was observed using increasing H_2_O_2_ concentrations in both mutant and control cells [24]. Therefore, we first performed MTT assays with different H_2_O_2_ concentrations to find the minimal concentration in which significant differences in viability were observed between *DJ-1* mutant and control cells (Fig. S3). Subsequently, enzymatic activities were determined under OS conditions induced with 50 µM H_2_O_2_, a concentration in which enough cells could be harvested. Our results showed that the activities of the key regulatory glycolytic enzymes Hk, Pfk and Pk, and especially of Eno were significantly increased in *DJ-1*-deficient cells compared to controls (Fig. 2). To exclude the possibility that variations in enzymatic activities between mutant and control cells were reflecting differences in gene expression levels, we performed RT-qPCR assays. Searches in FlyBase using the *Drosophila Eno* (CG17654), *Pfk* (CG4001), *Hex-A* (CG3001) and *PyK* (CG7070) genes as query revealed the existence of different orthologs in humans encoding glycolytic isozymes whose activities could have been measured in our assays. Among them, we selected for the RT-qPCR analysis those genes that are significantly expressed in SH-SY5Y cells according to the Human Protein Atlas [32,33]. These genes were: *PFKP* and *PFKM* (*Pfk* orthologs), *ENO1* and *ENO2* (*Eno* orthologs), *HK1* (*Hex-A* ortholog), and *PKM* (*PyK* ortholog). No significant differences in expression levels between *DJ-1*-deficient and control cells were found in the genes analyzed (Fig. S4). Therefore, the results obtained in *DJ-1*-deficient SH-SY5Y neuroblastoma cells are consistent with those obtained in *DJ-1β* mutant flies and suggest that loss of *DJ-1* function in both PD models produces an increase in the glycolytic rate. These results also confirm the translatability of the results obtained in *DJ-1β* mutant flies as reported in previous studies [24].

**Figure 2.**
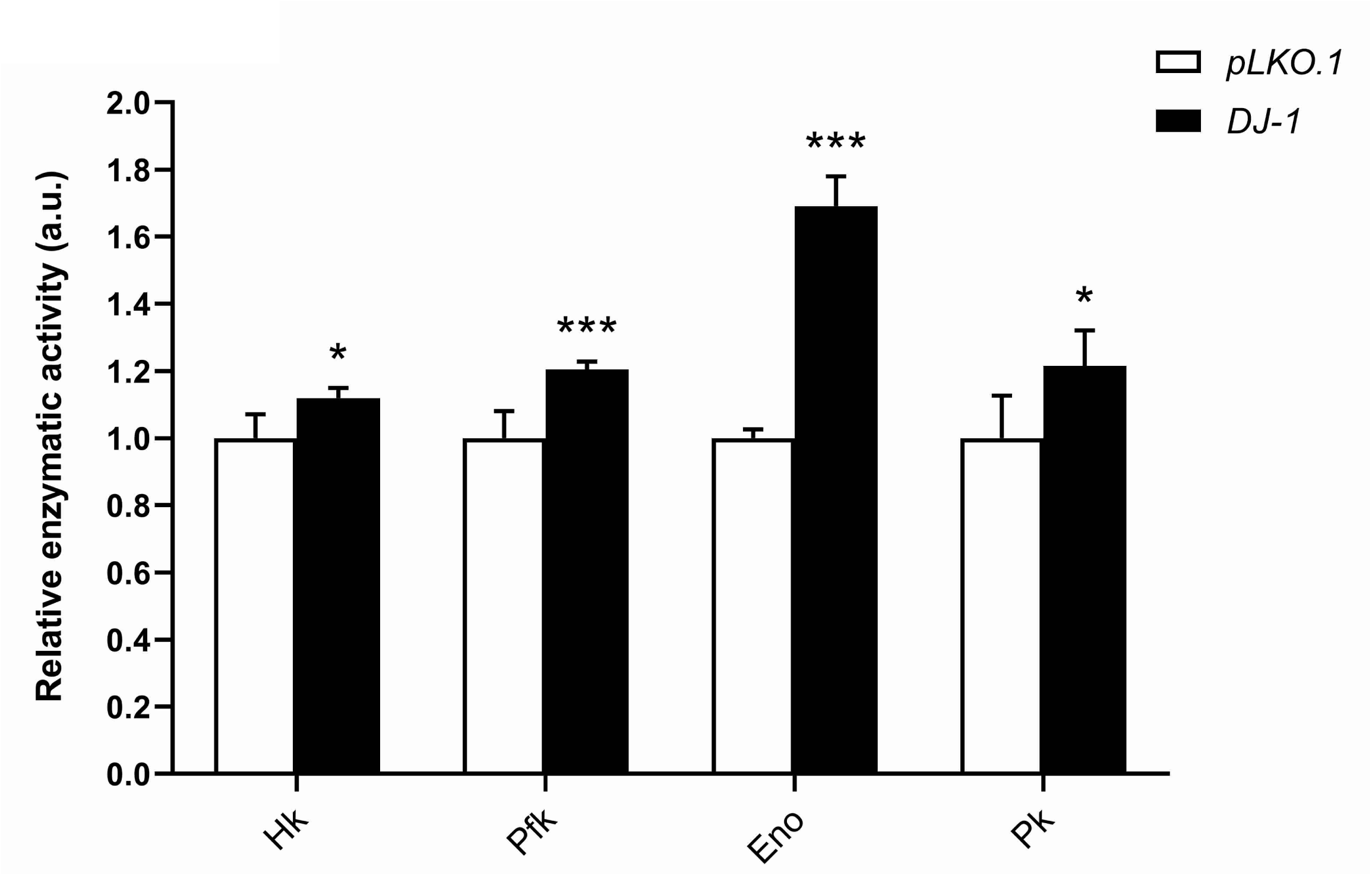
Activity of glycolytic enzymes in *DJ-1*mutant human cells. Hexokinase (Hk), phosphofructokinase (Pfk), enolase (Eno) and pyruvate kinase (Pk) enzymatic activities in *DJ-1* mutant cells. Results were relativized to data obtained in *pLKO.1* control cells and are expressed as arbitrary units (a.u). Error bars show s.d. from three replicates and three independent experiments (*, *P* < 0.05; ***, *P* < 0.001).

### Meclizine attenuates OS-induced citotoxicity and enhances glycolysis in *DJ-1*-deficient human cells

Recent studies performed in several genetic and chemical PD models have demonstrated that enhancing glycolysis has disease-modifying effects by different mechanisms such as increasing ATP content and dopamine levels, decreasing α-synuclein aggregates or controlling ROS production [20–22]. Hence, we decided to test whether a further increase in the glycolytic rate could suppress PD-related phenotypes in *DJ-1*-deficient human cells as well as in *DJ-1β* mutant flies. Meclizine (MEC) is an antihistaminic agent used to treat disequilibrium, vertigo and nauseas. It has been also demonstrated to protect against neuronal cell death in stroke, Huntington’s disease and PD models as well as to enhance glycolysis [34–37]. The mechanism by which MEC enhances glycolysis is by increasing the levels of 6-phosphofructo-2-kinase/fructose-2,6-biphosphatase 3 (*PFKFB3*) expression, which activates Pfk [36]. Thus, we first determined whether MEC was able to exert a protective effect on OS-induced cell death in *DJ-1-*deficient cells using an MTT assay. To this end, mutant cells were pretreated with five concentrations of MEC in a range 1-50 μM. OS was induced with 100 µM H_2_O_2_, a concentration in which survival of *DJ-1-*deficient cells was highly affected compared to control cells (Fig. S3). Our results showed that pretreatmens with MEC were able to attenuate cell death caused by H_2_O_2_ and that this effect was dose-dependent, being 50 µM the most effective MEC concentration in suppressing that phenotype (Fig. 3A). Subsequently, we performed enzymatic assays in cells pretreated with 50 µM MEC to determine whether this compound could affect the activity of any of the four glycolytic enzymes previously studied in untreated *DJ-1* mutant cells. Our results showed that MEC was able to slightly increase Pfk activity, supporting previous studies [36]. Surprisingly, we also found that Hk activity was robustly increased. No modifications in Eno or Pk activities were observed (Fig. 3B).

**Figure 3.**
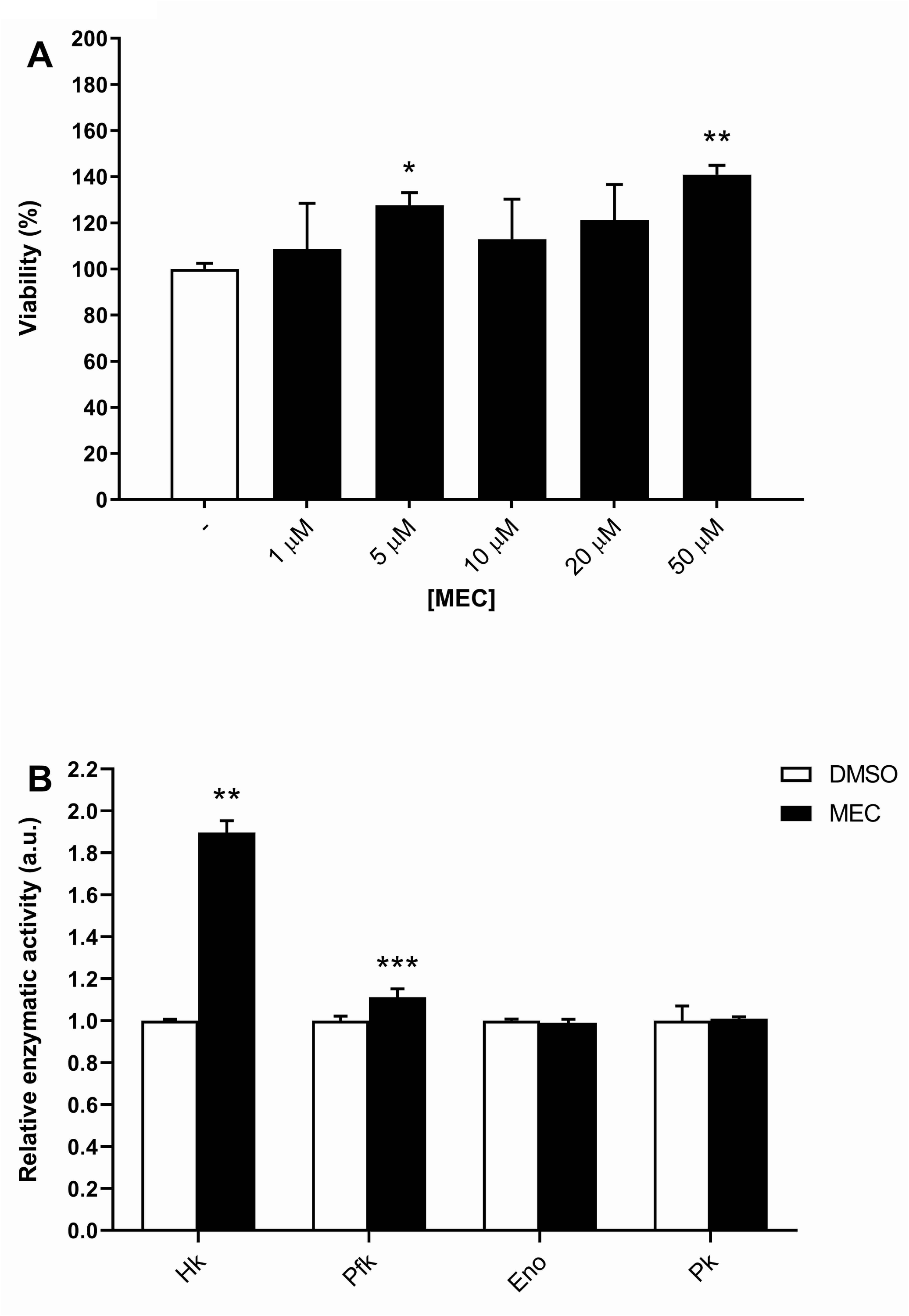
Effect of meclizine in viability and enzymatic activities of *DJ-1* mutant cells. (A) Viability of *DJ-1* mutant cells was measured by MTT assays in presence of OS (induced with 100 μM H_2_O_2_). Cells were either cultured in vehicle medium (0.1% DMSO) or treated with different concentrations of meclizine (MEC). Results were relativized to data obtained from *DJ-1* mutant cells cultured in vehicle (-). (B) Activity of glycolytic enzymes in *DJ-1* mutant cells treated with 50 µM MEC. Results were relativized to data from untreated *DJ-1* mutant cells cultured in vehicle (DMSO). In all cases, error bars show s.d. from three replicates and three independent experiments (*, *P* < 0.05; **, *P* < 0.01; ***, *P* < 0.001).

### Dimethyl fumarate attenuates OS-induced citotoxicity and enhances glycolysis in *DJ-1*-deficient human cells

Dimethyl fumarate (DMF) is a methyl ester of fumaric acid that is widely used to treat multiple sclerosis, psoriasis, cancer and other diseases [38–40]. It activates cellular antioxidant signaling pathways and may promote myelin preservation. However, it is still unclear what mechanisms may underlie this neuroprotection. It seems that at low concentrations DMF exerts an Nrf2-mediated antioxidant, neuroprotective and immunomodulator effect [41,42], which is independent of DJ-1 in primary neural cells and tissues [43]. However, high concentrations of DMF can cause OS and subsequently cytotoxicity in several cancer cells via Nrf2 depletion [44]. Several studies have demonstrated that DMF attenuates neurodegeneration in different PD models [41,45,46]. Interestingly, there are evidences suggesting the existence of a link between DMF and glycolysis. It has been shown that this compound appears to increase the glycolytic rate [42]. However, high DMF concentrations can downregulate glycolysis by inactivation of glyceraldehyde-3-phosphate dehydrogenase (GAPDH), a rate-limiting enzyme in the glycolytic pathway [47,48]. Considering this, we decided to test the effect of DMF in *DJ-1*-deficient cells. First, we determined whether DMF was able to exert a protective effect against OS-induced cell death. To this end, mutant cells were pretreated with five concentrations of DMF in a range 1-50 μM. Our results showed that pretreatmens with all DMF concentrations were able to attenuate OS-induced cell death in *DJ-1-*deficient cells, being 1 µM the most effective concentration in suppressing that phenotype (Fig. 4A). This is consistent with previous results showing that DMF has a cytoprotective role at low concentrations [44]. Subsequently, we performed enzymatic assays in cells pretreated with 1 µM DMF to determine whether this compound could affect the activity of any of the four glycolytic enzymes previously studied in untreated *DJ-1* mutant cells. Our results showed that cells pretreated with 1 µM DMF exhibited a significant increase in Pfk and Pk activities compared to cells treated with vehicle, although no differences in Hk and Eno activities were observed (Fig. 4B).

**Figure 4.**
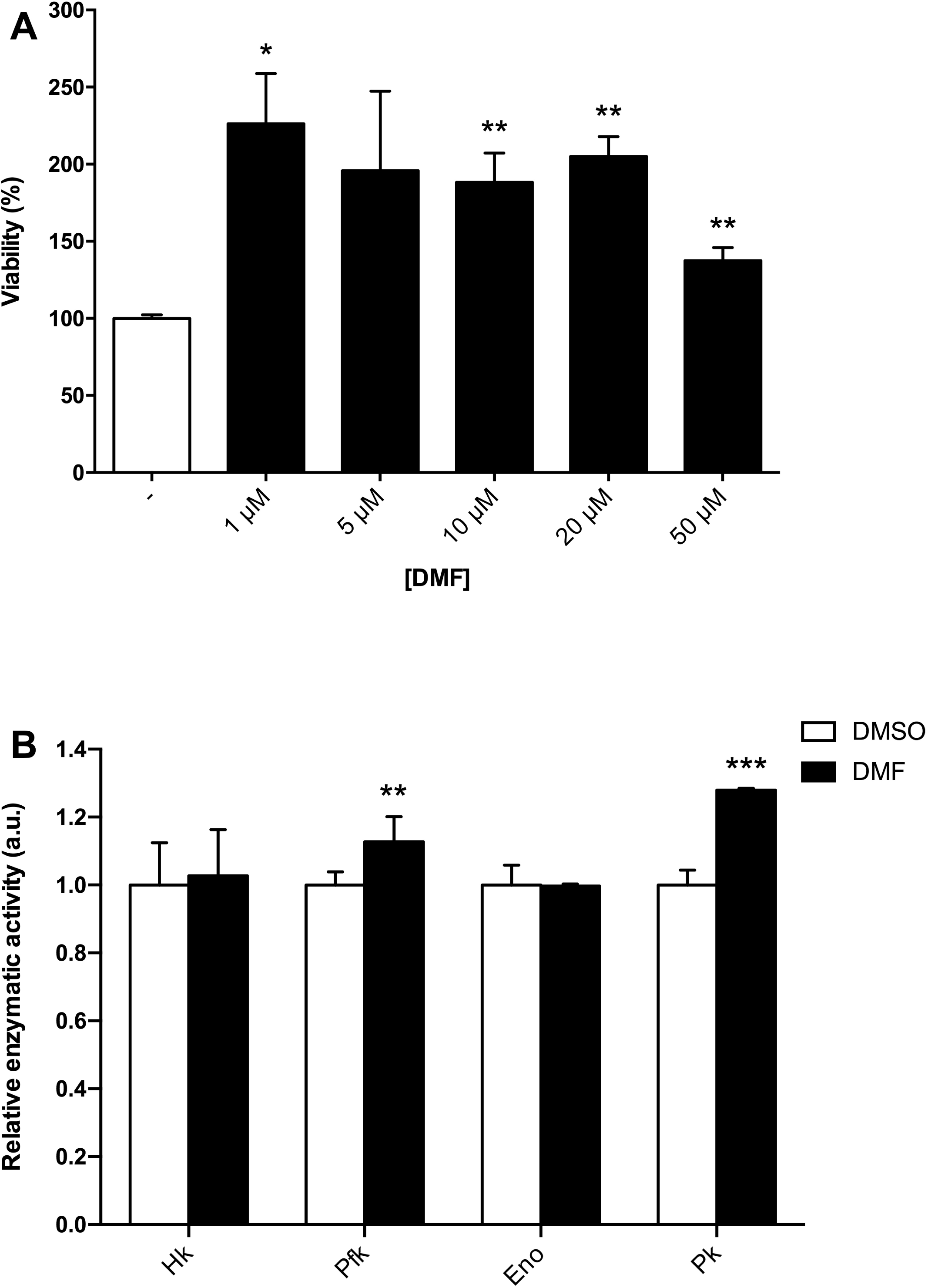
Effect of dimethyl fumarate in viability and enzymatic activities of *DJ-1* mutant cells. (A) Viability of *DJ-1* mutant cells was measured by MTT assays in presence of OS (induced with 100 μM H_2_O_2_). Cells were either cultured in vehicle medium (0.1% DMSO) or treated with different concentrations of dimethyl fumarate (DMF). Results were relativized to data obtained from *DJ-1* mutant cells cultured in vehicle (-). (B) Activity of glycolytic enzymes in *DJ-1* mutant cells treated with 1 µM DMF. Results were relativized to data from untreated *DJ-1* mutant cells cultured in vehicle (DMSO). In all cases, error bars show s.d. from three replicates and three independent experiments (*, *P* < 0.05; **, *P* < 0.01; ***, *P* < 0.001).

### Meclizine and dimethyl fumarate suppress phenotypes and enhance glycolysis in *DJ-1β* mutant flies by increasing Pfk activity

Once determined the effectiveness of MEC and DMF in *DJ-1-*deficient cells, we tested their effect in phenotypes displayed by PD model flies. *Drosophila* offers a more convenient way to evaluate the therapeutic potential of compounds since they can be tested in a whole organism context [49]. It was previously reported that *DJ-1β* mutants exhibited impaired motor performance, high ROS production as well as elevated protein carbonylation levels [11–13], phenotypes that were shown to be suppressed by treatments with potential therapeutic compounds [13,24]. To determine whether MEC and DMF were able to attenuate these phenotypes, *DJ-1β* mutants were cultured in medium supplemented with either MEC or DMF during development and flies were collected 5 days after eclosion. Climbing assays showed a significant improvement of motor performance in 5-day-old *DJ-1β* mutant flies treated with either MEC or DMF (Fig. 5A). We also evaluated H_2_O_2_ production (a component of the total ROS pool) and protein carbonylation levels in *DJ-1β* mutants treated with MEC or DMF. Our results showed that H_2_O_2_ production and, consistently, protein carbonylation levels were significantly reduced in *DJ-1β* mutants supplemented with the compounds (Fig. 5B-C). Finally, to investigate whether these compounds could affect the activity of glycolytic enzymes in *DJ-1β* mutant flies, we performed enzymatic activity assays in 15-day-old PD model flies treated with either MEC or DMF. Our results showed that both compounds were able to increase Pfk activity in *DJ-1β* mutants. However, no differences in Hex-A, Eno and PyK activities were observed (Fig. 6).

**Figure 5.**
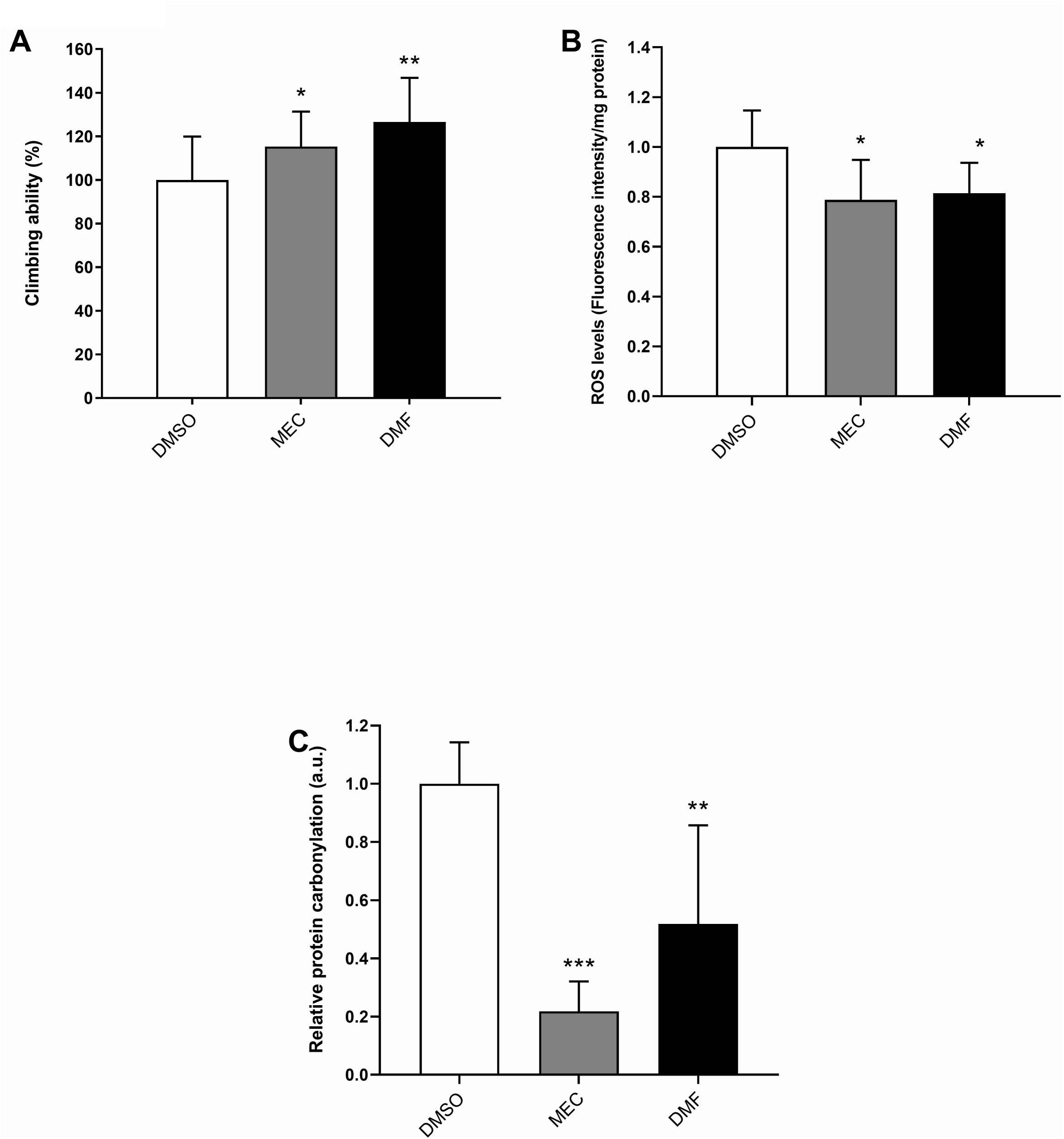
Analysis of locomotor performance, H_2_O_2_ levels and protein carbonylation levels in *DJ-1β* mutant flies treated with meclizine or dimethyl fumarate. (A) Motor performance of *DJ-1β* mutants treated with 50 µM MEC or 7 µM DMF was analyzed using climbing assays. (B) Levels of H_2_O_2_ production in *DJ-1β* mutant flies treated with 50 µM MEC or 7 µM DMF were analyzed by using the *Amplex H*_*2*_*O*_*2*_ *Red Kit* (Invitrogen). Data were expressed as arbitrary units (a.u.) per mg of proteins. (C) Protein carbonylation levels in *DJ-1β* mutant flies treated with 50 µM MEC or 7 µM DMF were analyzed by absorbance. In all cases, results were referred to data obtained in flies cultured in vehicle medium (DMSO). Error bars show s.d. from three replicates and three independent experiments (*, *P* < 0.05; **, *P* < 0.01; ***, *P* < 0.001).

**Figure 6.**
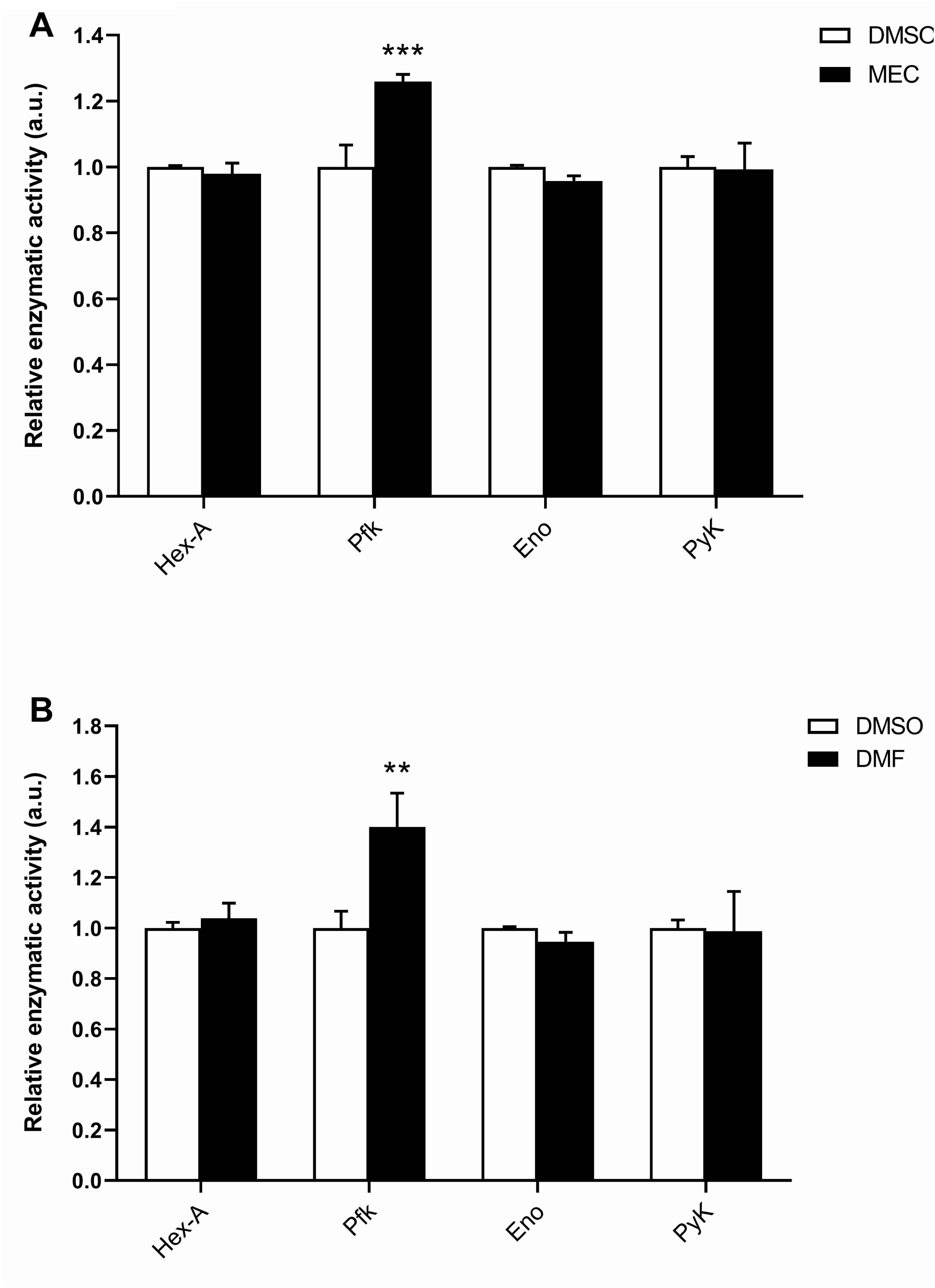
Effect of meclizine and dimethyl fumarate in enzymatic activities of *DJ-1β* mutant flies. Activity of glycolytic enzymes in *DJ-1β* mutant flies treated with (A) 50 µM MEC or (B) 7 µM DMF. Results were relativized to data obtained in flies cultured in vehicle medium (DMSO). Error bars show s.d. from three replicates and three independent experiments (**, *P* < 0.01; ***, *P* < 0.001).

Taken together, our results indicate that both MEC and DMF are able to suppress phenotypes in *DJ-1-*deficient cells and *DJ-1β* mutant flies. They also support previous findings indicating that MEC is able to increase the activity of Pfk, which in turn enhances glycolysis [36]. Increasing glycolysis may lead to a decrease in ROS production. Therefore, the improvement of motor performance observed in *DJ-1β* mutant flies after MEC and DMF supplementation is probably mediated by a reduction of ROS and protein carbonylation levels. Previous studies already showed that both appear to have a causative role in motor impairments [24]. In summary, our results support recent findings indicating that an enhancement of glycolysis may exert a beneficial effect in PD and could be a potential therapeutic target to slow or prevent the disease [20–22].

## DISCUSSION

In this work, we demonstrate for the first time that the activity of several glycolytic enzymes is increased in *Drosophila* and human cell PD models based on *DJ-1* inactivation. Our results also show that FDA-approved compounds such as MEC and DMF, which have different clinical applications, are able to attenuate PD-related phenotypes in both models. Moreover, we found that they could exert their beneficial effect by increasing glycolysis through the activation of key glycolytic enzymes. Therefore, an enhancement of the glycolytic pathway could be a protective mechanism in *DJ-1*-associated PD thus decreasing ROS production and restoring ATP levels caused by mitochondrial dysfunction.

Many neurodegenerative diseases such as PD or Alzheimer’s disease (AD) are usually characterized by mitochondrial impairment as well as increased ROS levels [50]. ROS significantly contribute to the deterioration of neuronal cells via modulating the function of biomolecules. Among them, proteins can be directly damaged by ROS through addition of carbonyl groups in particular amino acid side chains [14]. Although protein carbonylation is a posttranslational modification often associated to protein dysfunction, several evidences indicate that oxidized proteins can also acquire a gain of their function [14,15]. It has been reported that increased oxidation levels of proteins like glutathione peroxidase-1 in human umbilical vein endothelial cells [51], cathepsin D in Down syndrome [52] or superoxide dismutase 1 in amyotrophic lateral sclerosis [53] resulted in an increase in their enzymatic activity. In this work, we demonstrate that the glycolytic enzymes Pfk and Eno present a significantly increased activity in *DJ-1β* mutants compared to control flies, which may lead to an enhancement of the glycolytic pathway. Both enzymes were identified in a redox proteomics assay among a number of proteins highly oxidized in those mutants (unpublished data). Thus, changes in their activity could be the consequence of their increased oxidation levels in PD model flies. Pfk is the main regulatory enzyme in glycolysis and catalyzes the Mg-ATP-dependent transformation of fructose-6-phosphate into fructose-1,6-biphosphate. It is highly regulated by different metabolites such as ADP; a low ratio of ATP to ADP will activate the enzyme as well as glycolysis [29]. Since *DJ-1*-deficiency has been shown to decrease ATP levels [17,54], Pfk activity in *DJ-1β* mutant flies and *DJ-1-*deficient human cells may be increased in response to this signal to restore ATP levels and control ROS production. It is likely that the oxidative modification of Pfk in PD model flies could be affecting critical amino acid residues, either in the active site or in effector binding-sites, that may explain its increased activity [55]. Eno, which catalyzes the step in which 2-phosphoenolpyruvate (2PEP) is formed from 2-phosphoglycerate, appears to be one of the most consistently oxidatively modified proteins in brain of subjects with early-onset AD [56], which supports our redox proteomics results. However, it is unclear whether this is either due to its proximity to the many redox reactions occurring throughout the cell or to its structural susceptibility to oxidation. In fact, both possibilities appear reasonable since Eno is localized in different regions of the cell and possesses many active-site Lys and His residues that are extremely susceptible to carbonylation [57]. According to our results, Eno was also found to present higher carbonylation levels in brains of A30P α-synuclein transgenic PD model mice and a decreased activity when compared to wild-type mice [58]. This discrepancy is probably due to differences in the enzymatic assays used to measure Eno activity between both studies. However, an increase of Eno activity observed in *DJ-1β* mutant flies is consistent with the Pfk results, and points to an enhancement of glycolysis not only in *DJ-1β* mutant flies but also in *DJ-1-*deficient cells. Further analysis will be required to confirm whether both enzymatic activities are increased due to oxidative modification.

Glycolysis is a highly regulated multistep pathway that converts glucose into energy and substrates for other metabolic pathways [29]. Although Pfk is the most important control site in glycolysis as a first rate-limiting step, it is known that other enzymes such as PyK/Pk and Hex-A/Hk are involved in important regulatory steps of this pathway. Therefore, in this work we also analyzed the activity of such enzymes in PD model flies as well as in *DJ-1*-deficient human neuroblastoma cells. Our results showed that while PyK/Pk activity was increased in both models compared to controls, Hk activity was only increased in the cell PD model. PyK/Pk is the last regulatory enzyme of glycolysis, which catalyzes 2PEP transformation into pyruvate and is weakly regulated by ATP [29]. An increase of its activity in the *Drosophila* and human cell PD models may be induced by an accumulation of glycolytic metabolites due to the activation of previous steps in this pathway. Regarding Hk, this enzyme catalyzes the first step of the glycolytic pathway, which transforms glucose into glucose 6-phosphate (G6P), a branch point connecting glycolysis to other pathways such as the pentose phosphate pathway (PPP) [29]. PPP branches off glycolysis, utilizing the first glycolytic product G6P, and constitutes an essential pathway to produce NADPH, which is an important reducing agent essential for scavenging ROS and restore glutathione from its oxidized form [17,29]. Therefore, besides the obvious effects due to its glycolytic function, an increase of Hk activity in *DJ-1-*deficient human cells may play an antioxidant role by leading to G6P-derived NADPH production in response to high ROS levels [24]. Unexpectedly, Hex-A activity was not modified in PD model flies, in which high ROS levels were also detected [12,13]. Thus, different antioxidant mechanisms could be operating in this organism. Taken together, our results demonstrate that an increase in glycolysis due to an enhanced activity of glycolytic enzymes in PD models based on *DJ-1* deficiency could be a way to counteract decreased ATP levels and increased ROS production. It would be very interesting to determine whether these enzymes could directly suffer an oxidative modification that modifies their activity. Metabolic alterations have been previously described in studies performed in different PD models [16–18,59]. Indeed, PD is associated with mitochondrial dysfunction and, consequently, impaired oxidative phosphorylation (OXPHOS). Accordingly, loss of *DJ-1* function is known to inhibit complex I of the electronic transport chain (ETC) in the mitochondria, thus disrupting OXPHOS, decreasing ATP levels and increasing ROS production [17,54]. Interestingly, an enhancement of the glycolytic rate was also observed in MEFs obtained from *DJ-1* mutant mice [18], which is consistent with the results obtained in the present study in PD model flies and in *DJ-1*-deficient human cells. Therefore, enhancing glycolysis could constitute a compensatory mechanism to restore ATP levels and reduce ROS production in PD models based on *DJ-1* inactivation, as mentioned above. Similar results have been obtained in other fPD and iPD models. For example, it was shown that loss of *Pink1* function causes a metabolic reprogramming that promotes glucose uptake and glycolysis in MEFs, in mouse cortical neurons and also *in vivo* [19]. The authors proposed that mitochondrial OXPHOS defects in *Pink*-deficient cells are responsible for the switch from mitochondrial oxidative to glycolytic metabolism. Increased glycolysis has been also observed in chemically-induced iPD cell models such as murine brain neuroblastoma N2A cells or C6 glioma cells treated with 1-methyl-4-phenyl-1,2,3,6-tetrahydropyridine (MPTP), an active neurotoxin used to mimic PD pathophysiology that inhibits complex I of the ETC [60–62]. It was demonstrated that enhancing glycolysis protected such cells from MPTP toxicity by partially restoring ATP levels [60–62].

Recent reports have shown that increasing glycolysis by treating different animal and cell models with potential therapeutic compounds was able to attenuate PD-related phenotypes [20–22]. For example, treatments with hydrogen sulfide, a gasotransmitter that modulates several nervous, cardiovascular, and inflammatory functions in the organism [63], was proved to prevent DA neuron loss in 6-hydroxydopamine-exposed rats through leptin upregulation [22]. Terazosin, an activator of the glycolytic enzyme phosphoglycerate kinase 1, was able to restore PD symptoms in toxin-induced and genetic PD models in mice, rats, flies and human iPSC-derived neurons [20]. In this work, we demonstrate that MEC and DMF supplementation can suppress phenotypes caused by *DJ-1* inactivation in flies and human neuroblastoma cells probably by enhancing glycolysis. MEC was already proved to play a neuroprotective role in PD models [36,37]. It was shown that MEC supplementation during hypoxic exposure prevents ATP depletion, preserves NADPH and glutathione stores, curbs ROS and attenuates mitochondrial clustering in dorsal root ganglion neurites (DRG) [37]. Besides, MEC protected against 2-hydroxydopamine-induced apoptosis and cell death in SH-SY5Y human cells as well as in rat primary cortical neurons [36]. This effect was exerted by increasing the activity of PFKFB3 that catalyzes the synthesis of fructose-2,6-bisphosphate, an allosteric activator of Pfk [36]. Consistently, we have found that MEC supplementation increases Pfk activity in PD model flies and in *DJ-1*-deficient human cells. However, our results also showed that it increases Hk activity in the cell PD model. As mentioned above, G6P is the Hk product but also the substrate of the PPP, an extremely important antioxidant pathway that restores NADPH levels [17,29]. Since it was demonstrated that MEC was able to preserve NADPH in DRG [37], our results suggest that MEC may exert an antioxidant function through Hk activation in *DJ-1-*deficient human cells. DMF is an ester of fumaric acid widely studied for its anti-inflammatory properties [48], and described as an activator of Nrf2 [64]. DMF has been reported to reduce neurodegeneration in several PD models [41,45,46]. Different studies have demonstrated that this compound is related with glucose metabolism. Ahuja et al [42] showed that DMF was able to increase glycolysis in MEFs. However, DMF succinates the glycolytic enzyme glyceraldehyde 3-phosphate dehydrogenase (GAPDH), modification that inactivates this protein thus reducing glycolysis [47]. Our results demonstrated that DMF supplementation increases Pfk activity in PD model flies and in *DJ-1*-deficient human cells, but it also increases Pk activity in the cell PD model. It has been reported that Nrf2 promotes the expression of *PFKFB3* [65]. Therefore, DMF may lead to an increase of Pfk activity through PFKFB3 [66]. Besides, previous studies have shown that treatment of oligodendrocytes with DMF induced changes in citric acid cycle intermediates. Among them, succinate and fumarate were significantly upregulated in treated cells compared to controls [67]. These intermediates could increase oxalacetate production, which may be used to synthetize 2PEP. Since 2PEP is the substrate of Pk, this could explain the increase of Pk activity in *DJ-1*-deficient human cells treated with DMF. It should be mentioned that DMF (Tecfidera®) is currently approved for the treatment of multiple sclerosis. Besides, MEC is the generic name for the prescription drug called Antivert, which is indicated for the treatment of vertigo. Thus, our results support further evaluation and potential repurposing of both MEC and DMF as modulators of energy metabolism for neuroprotection in PD.

In summary, we demonstrate that an enhancement of glycolysis may have beneficial consequences in PD models as recently published, probably by increasing OXPHOS, mitochondrial activity as well as ATP concentration [68]. Taken together, these results open new directions for potential disease-modifying therapy in PD targeting the glycolytic pathway [69].

## Abbreviations

DA: dopaminergic
DMF: dimethyl fumarate
DMSO: dimethyl sulfoxide
MPTP: 1-methyl-4-phenyl-1,2,3,6-tetrahydropyridine
MEC: meclizine
MEF: mouse embryonic fibroblast
OS: oxidative stress
OXPHOS: oxidative phosphorylation
PD: Parkinson’s disease
ROS: reactive oxygen species

## ACKNOWLEDGMENTS

We are grateful to Dr. Jongkyeong Chung, the Bloomington *Drosophila* Stock Center and the Vienna *Drosophila* Research Center for providing fly stocks.

## FUNDING

This work was supported by Generalitat Valenciana [grant number PROMETEO/2014/067 to N.P.] and the University of Valencia [grant number UV-INV-AE17-702300 to N.P.].

## Supplementary figures

**Figure S1.**
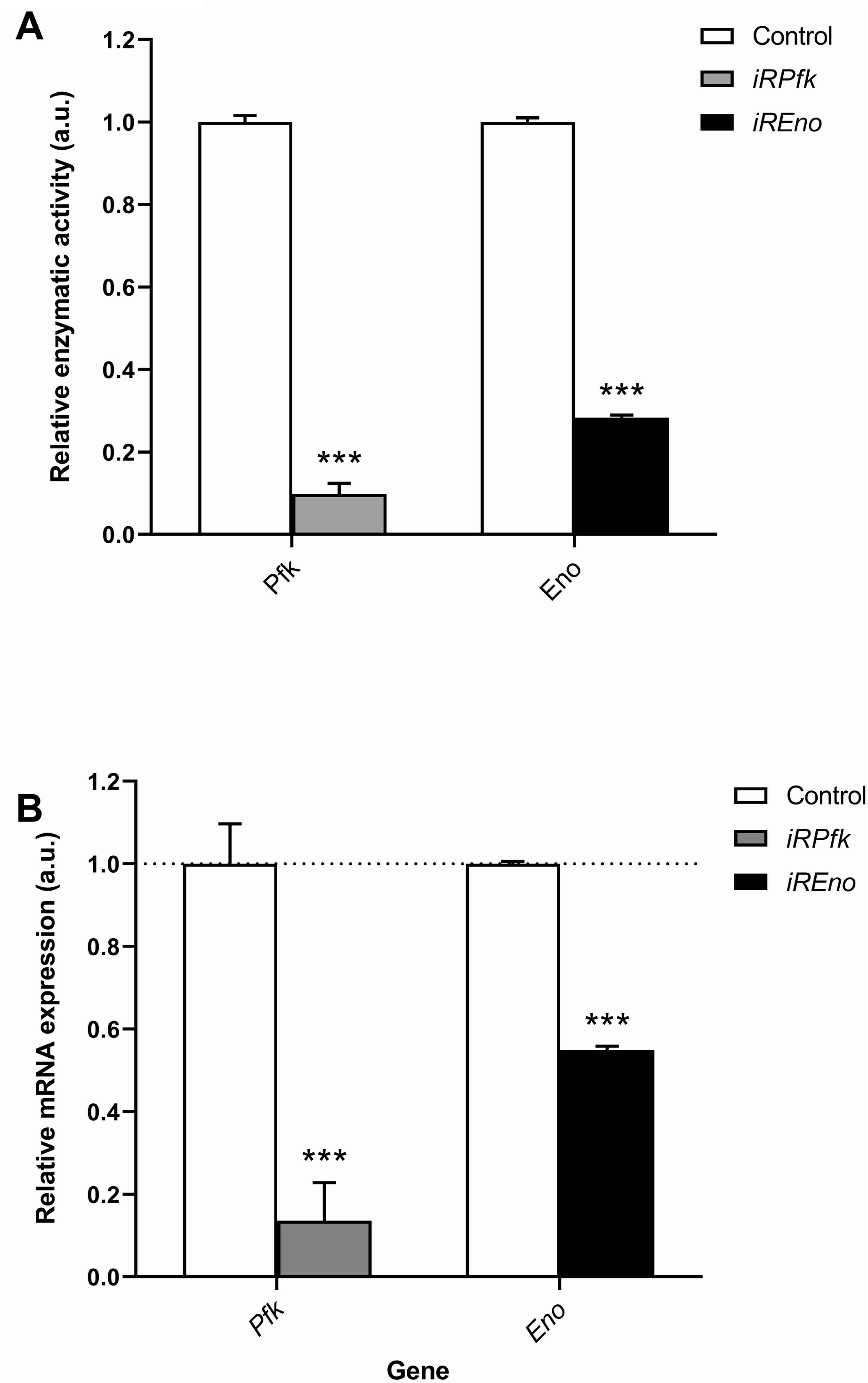
Analysis of enzymatic activities and gene expression levels in *iRPfk* and *iREno* mutant flies. (A) Pfk and Eno enzymatic activities were measured in 15-day-old *iRPfk* and *iREno* mutant flies. Results are relativized to data obtained from *arm-*GAL4/+ flies (control) and are expressed as arbitrary units (a.u.). Error bars show s.d. from three independent experiments in which three biological replicates were used. (B) Graphical representation of expression levels of *Pfk* and *Eno* genes quantified by RT-qPCR analysis in 15-day-old *iRPfk* and *iREno* flies. Results are referred to data obtained in *arm*-GAL4/+ flies (control). *tubulin* expression levels were measured and used as an internal control for RNA amount in each sample. Error bars show s.d. from four independent experiments (***, *P* < 0.001).

**Figure S2.**
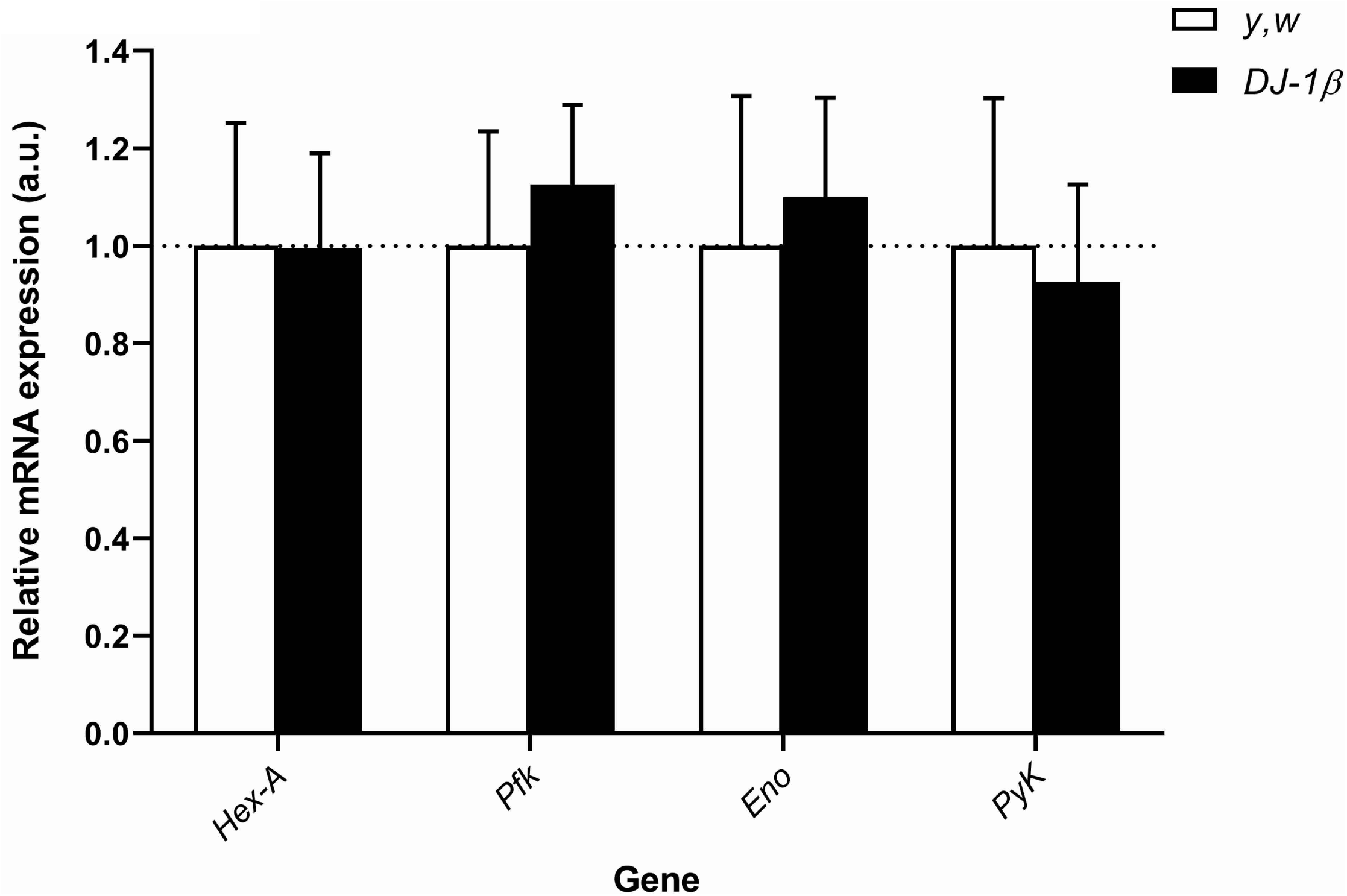
Analysis of gene expression levels in *DJ-1β* mutant flies. Graphical representation of expression levels of *Hex-A, Pfk, Eno*, and *PyK* genes quantified by RT-qPCR analysis in *DJ-1β* mutants. Results are referred to data obtained in *y,w* control flies and are expressed as arbitrary units (a.u.). *tubulin* expression levels were measured and used as an internal control for RNA amount in each sample. Error bars show s.d. from four independent experiments. No significant differences in gene expression levels were observed between *DJ-1β* mutants and control flies.

**Figure S3.**
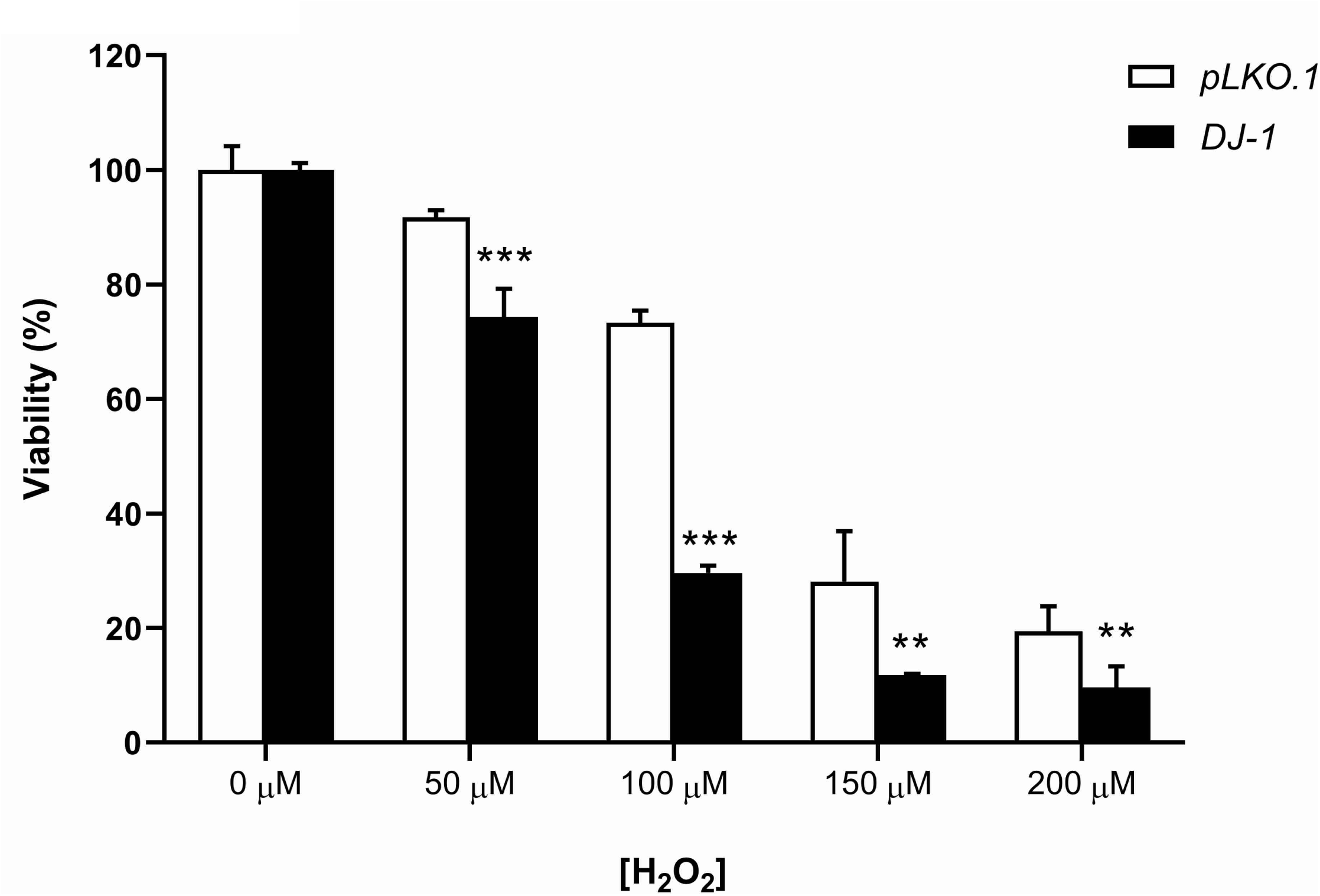
Viability assays in *pLKO.1* and *DJ-1-*deficient human neuroblastoma cells. Viability was measured by MTT in control and mutant cells cultured in absence and presence of OS, which was induced by different H_2_O_2_ concentrations (50, 100, 150 and 200 µM). Results were relativized to data obtained from *pLKO.1* control cells in absence of OS, and are expressed as arbitrary units (a.u.). Error bars show s.d. from three independent experiments in which three biological replicates were used (**, *P* < 0.01; ***, *P* < 0.001).

**Figure S4.**
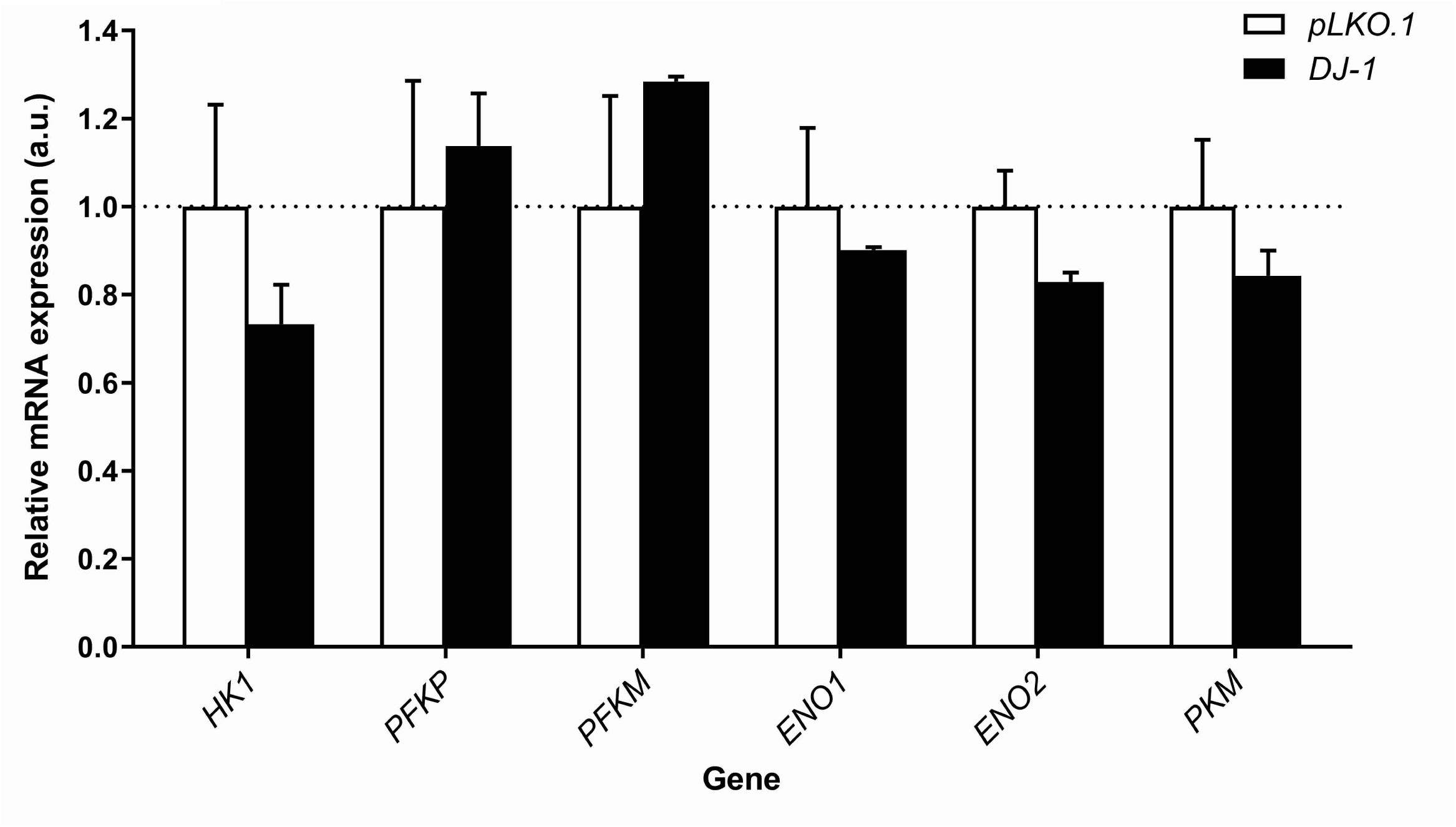
Analysis of gene expression levels in *DJ-1-*deficient human neuroblastoma cells. Graphical representation of expression levels of *HK1, PFKP, PFKM, ENO1, ENO2* and *PKM* genes quantified by RT-qPCR analysis in *DJ-1* mutant cells under OS condition induced with 100 μM H_2_O_2_. Results are referred to data obtained in *pLKO.1* control cells under the same OS condition than mutant cells, and are expressed as arbitrary units (a.u.). *tubulin* expression levels were measured and used as an internal control for RNA amount in each sample. No significant differences in gene expression levels were observed between *DJ-1* mutant and control cells. Error bars show s.d. from four independent experiments.

